# Antisense oligonucleotide therapy for SCN2A gain-of-function epilepsy

**DOI:** 10.1101/2020.09.09.289900

**Authors:** Melody Li, Nikola Jancovski, Paymaan Jafar-Nejad, Lisseth Estefania Burbano, Ben Rollo, Kay Richards, Lisa Drew, Alicia Sedo, Svenja Pachernegg, Armand Soriano, Linghan Jia, Todd Blackburn, Blaine Roberts, Alex Nemiroff, Kelley Dalby, Snezana Maljevic, Christopher Reid, Frank Rigo, Steven Petrou

## Abstract

The clinical spectrum associated with SCN2A *de novo* mutations (DNMs) continues to expand and includes autism spectrum disorder with or without seizures, in addition to early and late seizure onset developmental and epileptic encephalopathies (DEEs). Recent biophysical studies on SCN2A variants suggest that the majority of early seizure onset DEE DNMs cause gain of function. Gain of function in SCN2A, the principal sodium channel of excitatory pyramidal neurons, would result in heightened neuronal activity and is likely to underlie the pathology seen in early seizure onset DEE patients. Supratherapeutic dosing of the non-selective sodium channel blocker phenytoin, is effective in controlling seizures in these patients but does not impact neurodevelopment, raising the idea that more profound and specific reduction in SCN2A function could significantly improve clinical outcome. To test the potential therapeutic benefit of reducing SCN2A in early seizure onset DEE we centrally administered an antisense oligonucleotide (ASO) targeting mouse Scn2a (Scn2a ASO) to a mouse model of human SCN2A early seizure onset DEE. Mice were genetically engineered to harbour the human equivalent SCN2A p.R1882Q mutation (Q/+), one of the most recurrent mutations in early seizure onset DEE. Q/+ mice presented with spontaneous seizures at postnatal day (P) 1 and did not survive beyond P30. Intracerebroventricular Scn2a ASO administration into Q/+ mice between P1-2 (that reduced Scn2a mRNA levels by 50%) significantly extended lifespan and markedly reduced spontaneous seizures occurrence. Across a range of cognitive and motor behavioural tests, Scn2a ASO treated Q/+ mice were largely indistinguishable from wildtype (+/+) mice. Further improvements in survival and behaviour were seen by adjustment of dosing regimens during development. Scn2a ASO efficacy was also evident at the cellular level. Whole cell patch clamp recording showed that Scn2a ASO administration reversed changes in neuronal excitability in layer 2/3 pyramidal neurons of Q/+ mice to levels seen in +/+ mice. Safety was assessed in +/+ mice and showed a developmental stage dependent tolerability and a favourable therapeutic index. This study suggests that a human SCN2A gapmer ASO could profoundly and safely impact early seizure onset DEE patients and heralds a new era of precision therapy in neurodevelopmental disorders.

## Introduction

Developmental and epileptic encephalopathies (DEEs) are devastating neurological disorders presenting during infancy and early childhood ^1^. The affected children have ongoing, refractory seizures in addition to profound global developmental delay, intellectual disability and movement disorders^1^. The prognosis of these patients is poor, marked by progressive disability and increased risk of early death ^1^. The limited treatment options, presence of comorbidities and need of long term supported care represent an important burden for affected patients, caregivers, and the health services.

Only a decade ago little was known about DEE aetiologies, but the employment of next generation sequencing has identified single gene *de novo* mutations (DNMs) as the main cause ^2^. The number of identified DEE genes is constantly increasing with those encoding ion channels as the pivotal brain excitability regulators being the most represented ^2-4^. Identification of genetic causes of DEE has allowed the mechanistic basis of the disease to be probed in a way not previously possible and has laid the foundation for the development of precision medicine therapeutic approaches. This is especially important as the current therapies focus on the symptomatic control of seizures while the developmental aspects of disease remain untended.

Parallel to the discovery of genetic causes of DEE, several major technological advances have reiterated the possibility of establishing effective gene therapies for these disorders. Especially, a recent approval of antisense oligonucleotides (ASOs) as a treatment for spinal muscular atrophy, Huntington’s disease and Duchenne Muscular Dystrophy showed that RNA targeted therapies can be a viable therapeutic strategy for these devastating neurogenetic disorders^5-7^. ASOs are single-stranded oligonucleotides that are designed to specifically target the RNA of interest through Watson-Crick base pairing which can result in RNase H1 mediated mRNA degradation, modulation of pre-mRNA splicing or regulation of translational efficiency ^8,9^.

SCN2A encodes the α subunit of the voltage-gated sodium channel (Na_v_1.2) and is involved in the initiation and conduction of action potentials, particularly during early post-natal development when expression is higher than other voltage-gated sodium channel isoforms ^10,11^. This gene has emerged as the most commonly mutated single gene causing neurodevelopmental disorders ^12,13^. The phenotypic spectrum associated with SCN2A DNMs includes schizophrenia, autism spectrum disorder, intellectual disability and a myriad of epileptic syndromes including DEE ^12-16^. Case history studies of patients with SCN2A DEE have identified two major clinical presentations based on the age of seizure onset and disease severity: early onset is characterised by seizures starting within the first 3 months of life and late onset seizures usually commence between 3 months and 4 years of age ^14^. Functional investigation of a number SCN2A DEE DNMs has shown that early seizure onset phenotype is caused by gain of function of SCN2A whereas the late seizure onset DEE is linked to a loss of function patho-mechanism ^12,17^. This fits well with the observation that patients with the early seizure onset respond better to antiepileptic drugs that non-selectively block sodium channel function, such as phenytoin ^14^. Moreover, the observation suggests that these patients could be amenable to gene therapy designed to reduce the SCN2A expression.

To corroborate this hypothesis, we engineered a syndrome-specific mouse model of SCN2A disease based on the human equivalent 5645G>A variant, that results in the protein change p.R1882Q. This is one of the most recurrent pathogenic variants for early seizure onset SCN2A DEE ^14^. Phenytoin partially improves seizure outcomes in some patients carrying p.R1882Q variant ^14^. However, within the therapeutic window of phenytoin, seizures are not fully resolved, and developmental impairment remains unaddressed. *In vitro* biophysical analysis confirmed the p.R1882Q variant causes larger peak and persistent sodium currents, with dynamic clamp simulations predicting a higher action potential firing frequency ^17^.

Here, an antisense oligonucleotide (ASO) specifically designed to down-regulate mouse Scn2a gene expression (Scn2a ASO) was delivered to our engineered mouse model (Q/+). The Q/+ mouse model develops spontaneous seizures as early as postnatal day one (P1) and suffers premature death recapitulating the severe symptoms observed in patients with early seizure onset DEE. We showed that Scn2a ASO mediated knockdown effectively prevented premature death, suppressed spontaneous seizures, and reversed pathological cellular excitability in the Q/+ mouse model. The Scn2a ASO treated Q/+ mice behaved similarly to wildtype (+/+) mice in motor and social function tests. Furthermore, down-regulation of Scn2a within the therapeutic range did not cause any adverse effects. Scn2a ASO is the first therapeutic entity to address both seizure and developmental impairment in SCN2A early seizure onset DEE, positioning ASO as a promising therapeutic strategy for SCN2A gain-of-function diseases.

## Results

### The Scn2a gain-of-function DEE mouse model

The SCN2A R1882Q variant is located at the C-terminal tail (Fig. 1a). Mirroring the clinical presentation of SCN2A p.R1882Q DEE, Q/+ mice have a severe, early seizure onset phenotype. Spontaneous tonic seizures were observed as early as P1, which later developed into tonic-clonic seizures with hindlimb extension (Fig. 1b). The earliest death recorded was P13 and the median survival was P18 (Fig. 1c). Body weight was measured on P21 and there was no significant difference between Q/+ and +/+ mice (Fig. 1d). No Q/+ mice survived beyond P30, which limited behavioural characterization. Phenytoin, the first-line therapy for early seizure onset SCN2A DEE patients ^14,17^, was administered to Q/+ mice via intra-peritoneal injection from P10 (20 mg/kg, daily). While phenytoin significantly extended median survival by three days when compared to vehicle administration, it failed to rescue the premature death phenotype (Extended data fig. 1).

**Figure 1.**
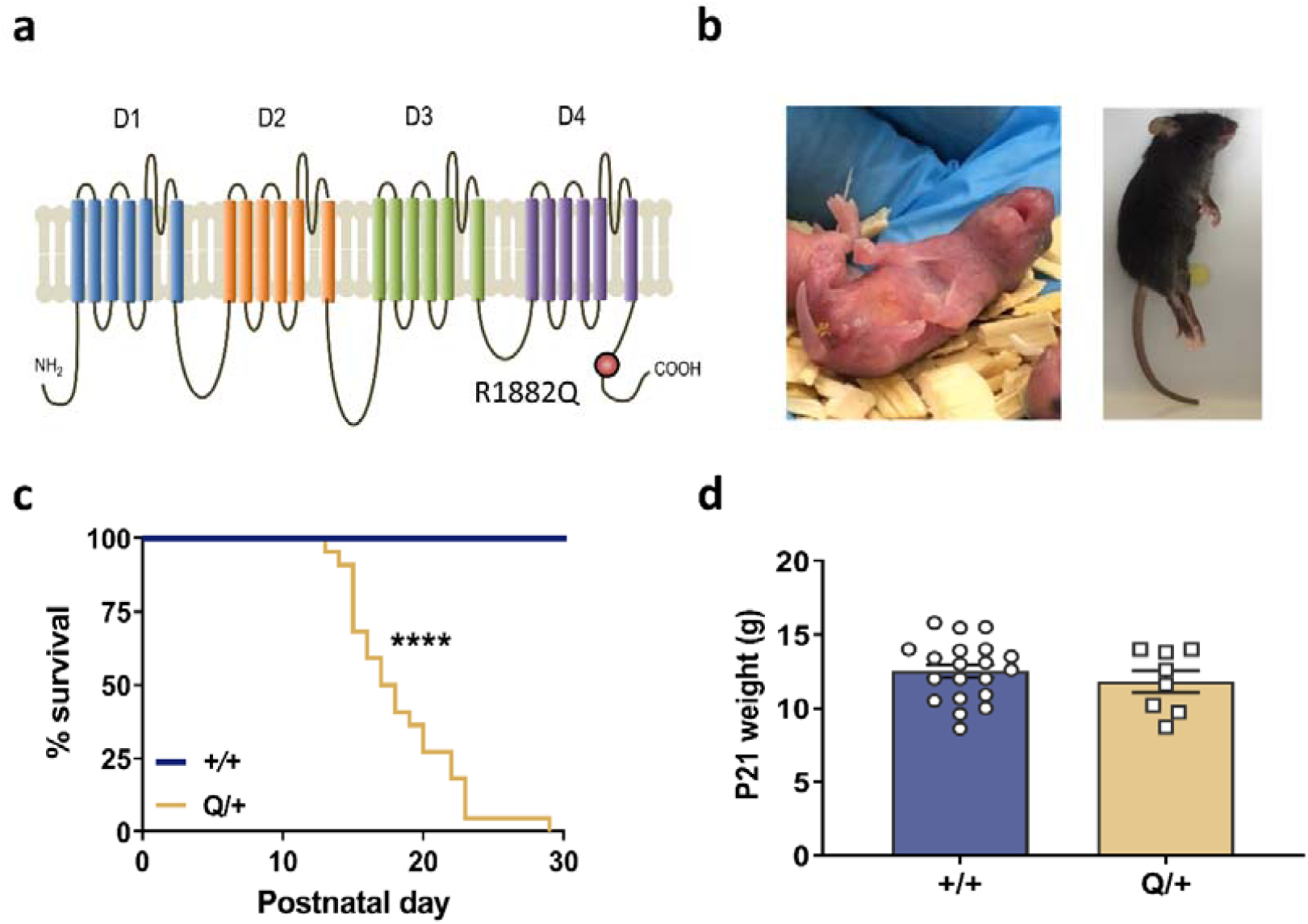
Disease phenotype of the Q/+ mouse model. **a.** Schematic presentation of SCN2A channel depicting four domains (D1-D4), each comprised of 6 transmembrane regions, and the intracellular N and C-terminus of the channel. R1882Q variant is predicted to affect the C-terminus of the channel. **b.** Images of Q/+ mice undergoing spontaneous seizure on P1 (left) and on P25 (right). **c.** Survival curves of Q/+ and +/+ mice. (+/+ n = 19, Q/+ n = 22). **** *P* < 0.0001, Log-rank test. **d.** Body weight measured on P21 (+/+ n = 20, Q/+ n = 8).

**Extended data figure 1.**
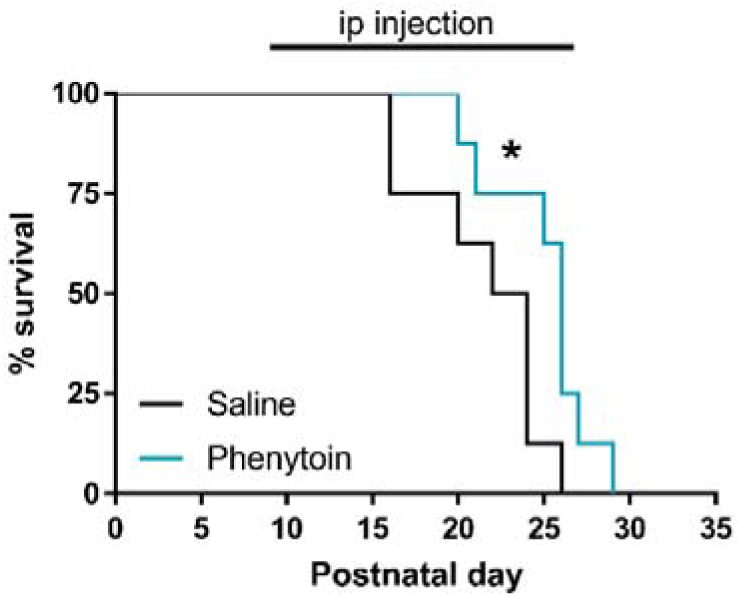
Phenytoin showed mild efficacy in the Q/+ mouse model. Survival curves of Q/+ mice ip injected daily with phenytoin (20 mg/kg) or saline from P10. Saline (n = 8), phenytoin (n = 8), * *P* < 0.05, Log rank test.

### Scn2a ASO mediated Scn2a reduction rescues premature death and seizure phenotype in Q/+ mice

To test whether ASOs targeting Scn2a could mitigate disease in the SCN2A early seizure onset DEE mouse model, we treated Q/+ mice with a single intracerebroventricular (ICV) injection of a non-targeting control ASO or Scn2a ASO at P1. Using a pan-ASO antibody, we confirmed broad ASO distribution throughout the mouse brain (Extended data fig. 2).

**Extended data figure 2.**
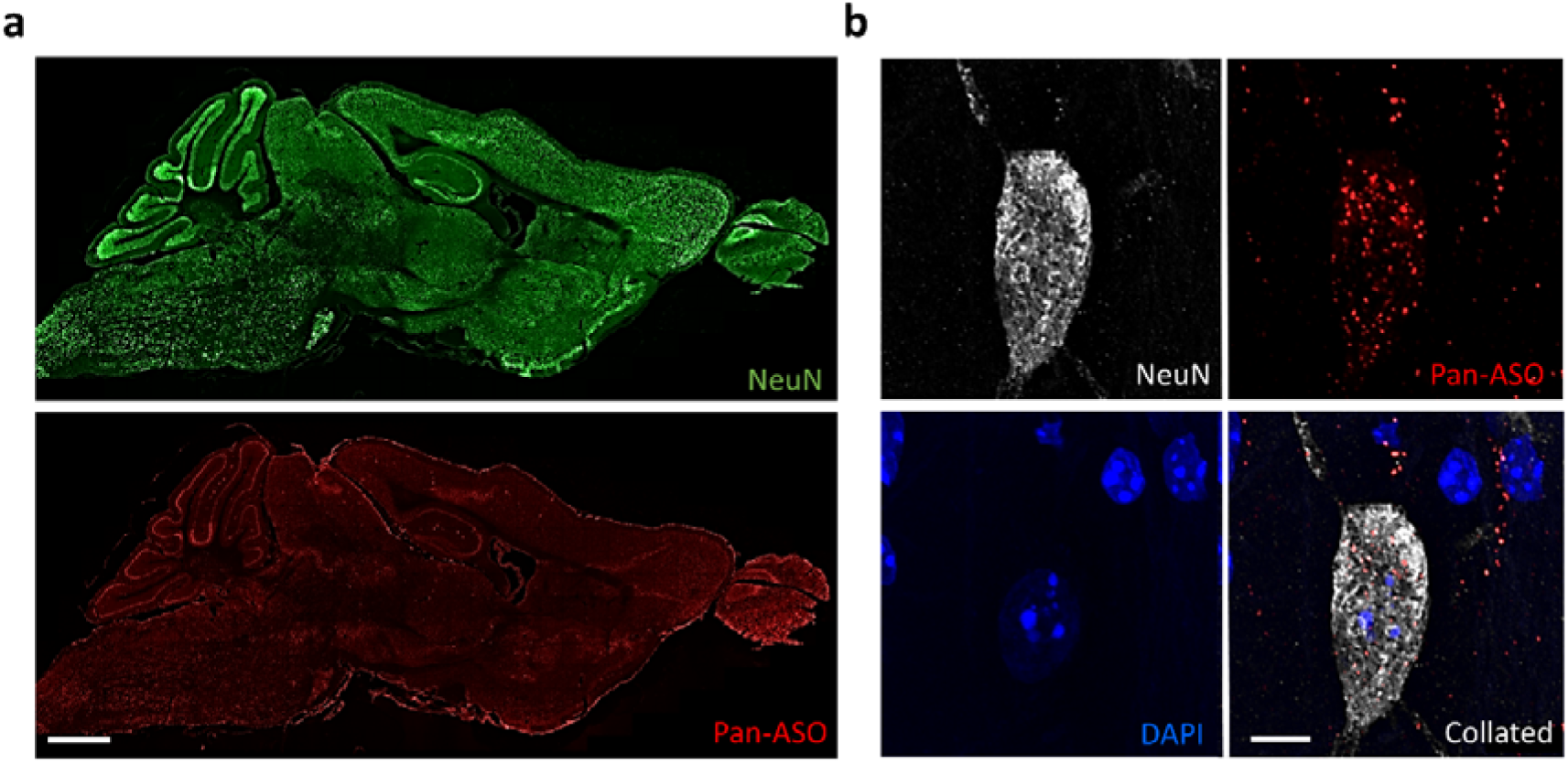
Distribution of Scn2a ASO in mouse brain. **a.** Sagittal brain section of a mouse icv injected with Scn2a ASO (500 µg) on P60. Brain was collected 12 days post-icv injection. Slice were stained with NeuN (top) and pan-ASO antibody (bottom). Scale bar represents 1 mm. b. Hippocampal neuron co-stained by NeuN, pan-ASO and DAPI. Scale bar represents 5 µm.

Furthermore, we confirmed that Scn2a ASO specifically reduces Scn2a mRNA and had no effect on the other voltage gated sodium channels (Fig. 2a). At the protein level, mass spectrometry determined a 3.8-fold reduction in cortical Scn2a protein levels in Scn2a ASO treated mice when compared to controls (Fig. 2b). Immunohistochemistry also confirmed Scn2a protein expression was reduced along the axonal initial segment (Fig. 2c). To determine the effective dose for 50 % (ED_50_) and 80 % (ED_80_) Scn2a mRNA reduction, mice were treated with Scn2a ASO at P1, P15, and P30 and Scn2a mRNA levels measured 2 weeks post treatment (Extended data fig. 3).

**Figure 2.**
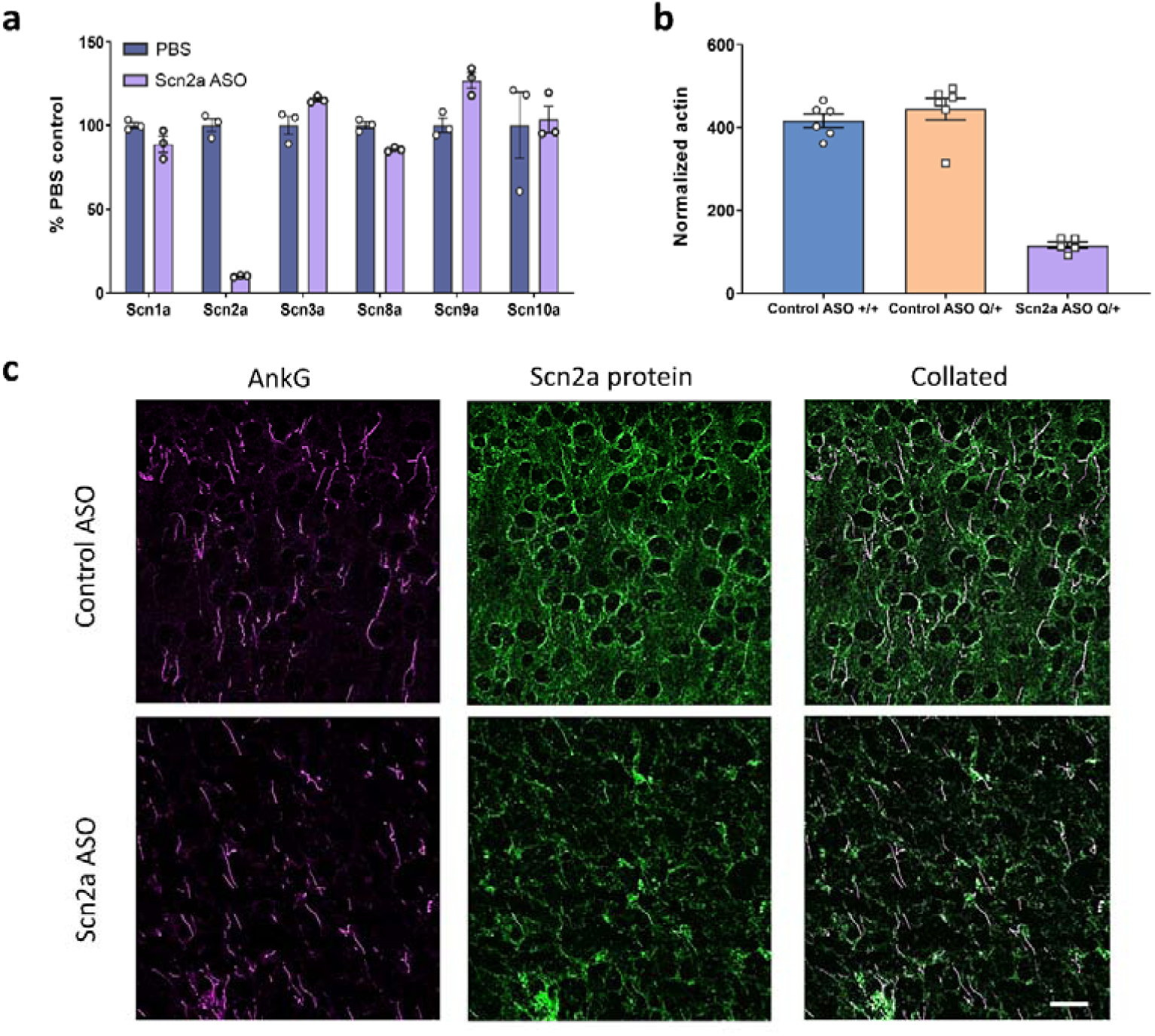
Scn2a mRNA and protein expression were reduced by Scn2a ASO. **a.** Percentage of Nav isoform mRNA remaining in the cortex. +/+ (P30-40) were icv injected with PBS or Scn2a ASO (500 µg). The mRNA level was measured 4 weeks post-injection, n = 3 per group. **b.** Level of Scn2a protein normalized to actin. **c.** Z-stack images of cortical slices labelled by AnkG (magenta) and Scn2a protein (green) from mice administered with control ASO (50 µg) or Scn2a ASO (10 µg) on P1. Scale bar indicates 20 µm.

**Extended data figure 3.**
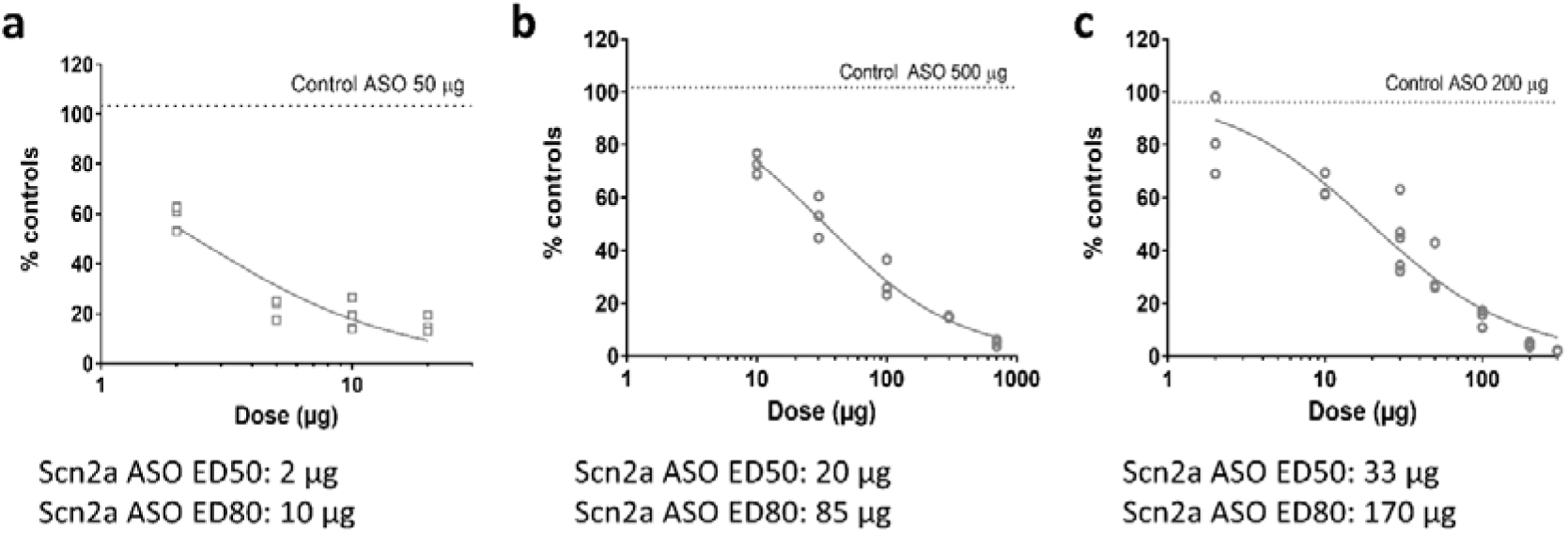
Scn2a ASO dose-dependently reduced Scn2a mRNA. Mice icv injected with control ASO or a range of Scn2a ASO dose on P1 (a), P15 (b) or P30 (c). The Scn2a mRNA level was assessed 14-15 days post-icv, and ED_50_ or ED_80_ were determined by Motulsky regression fit, N = 3 per dose.

Scn2a ASO administration at ED_50_ to Q/+ P1 mice increased the median survival to P47 and at ED_80_ median survival was improved even further to P87 (Fig. 3a). As expected, control ASO did not improve survival and median survival was P20 (Fig. 3a). To determine if the survival can be extended even further, we treated one day old Q/+ mice with Scn2a ASO at ED_50_, then at P27-P30 we administered a second dose of Scn2a ASO at ED_50_ or ED_80_ or control ASO. Q/+ mice that received a second dose of Scn2a ASO had significantly longer life span than those re-dosed with the control ASO (median survival control ASO: P86; Scn2a ASO at ED_50_: P145; Scn2a ASO at ED_80_: P206) (Fig. 3b). The repeated dosing later in development was able to maintain the therapeutic level required and was effective in acutely supressing disease re-emergence.

**Figure 3.**
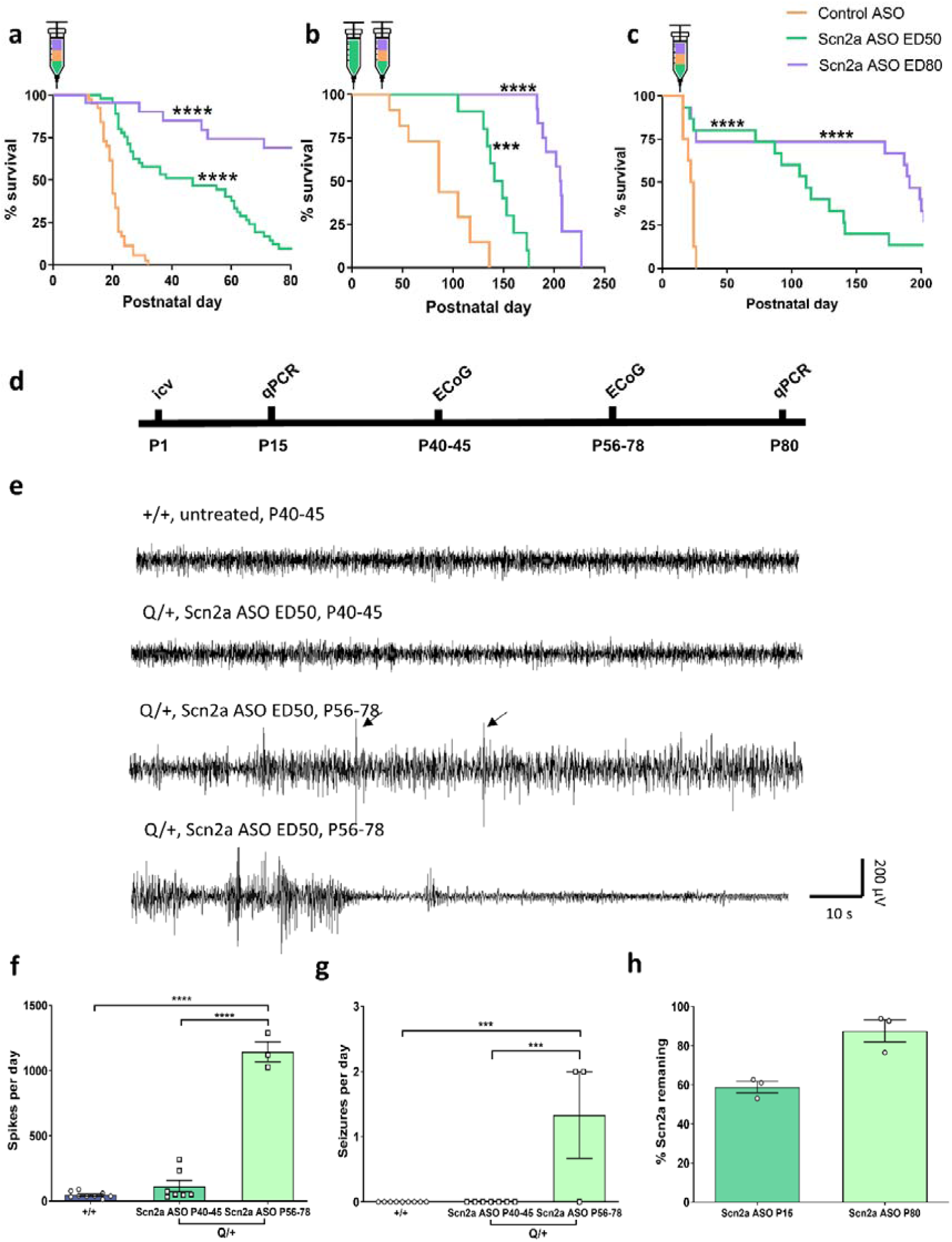
Scn2a ASO rescues the disease phenotype of Q/+ mice. **a.** Survival curves of Q/+ mice administered with Scn2a ASO or the negative control, control ASO on P1. Control ASO (50 µg, n = 39), Scn2a ASO ED_50_ (2 µg, n = 49) and Scn2a ASO ED_80_ (10 µg, n = 22). **b.** Survival curves of Q/+ mice administered with Scn2a ASO ED_50_ (2 µg) on P1, then re-dosed with control ASO (85 µg), Scn2a ASO ED_50_ (20 µg) or Scn2a ASO ED_80_ (85 µg) on P27-30. Control ASO (n = 11), Scn2a ASO ED_50_ (n = 15) and Scn2a ASO ED_80_ (n = 13). **c.** Survival curves of Q/+ mice administered with Scn2a ASO or control ASO on P14-16. Control ASO (170 µg, n = 8), Scn2a ASO ED_50_ (33 µg, n = 15) and Scn2a ASO ED_80_ (170 µg, n = 16). Syringes indicate time of icv injection. **d.** Timeline of ECoG and qPCR experiments. **e.** Representative ECoG traces recorded from treatment groups as per label. Arrows indicate inter-ictal spikes. Scale bar applies to all traces. **f.** Number of spikes during 24-hour ECoG recording. **g.** Number of electrical seizures during 24-hour ECoG recording. **h.** Percentage of Scn2a mRNA remaining in the cortex 14 days or 79 days after 890911 ED_50_ administration on P1. *** *P* < 0.005, **** *P* < 0.0001, Log-rank test (**a-c**), one-way ANOVA with Dunnett’s multiple comparison (**f-g**).

In practice, the ASO therapy would be delivered after genetic diagnosis and the DEE symptoms would have likely progressed before initiating ASO treatment. Therefore, it was critical to assess the efficacy of Scn2a ASO at later age of treatment initiation. Since Q/+ mice have spontaneous seizures at P1 and mortality was first observed at P13, Scn2a ASO was administered as a single bolus at P14-16. Lifespan of Q/+ mice were still significantly extended with the later therapeutic intervention. Scn2a ASO at ED_50_ and Scn2a ASO at ED_80_ extended median survival to P111 and P191 respectively (Fig. 3c). The median survival for control ASO was P23 (Fig. 3c). These data strongly indicate later therapeutic intervention with Scn2a ASO has compelling disease reversing capability.

Since Scn2a ASO treatment significantly extended survival, its seizure modifying ability was then ascertained. After Scn2a ASO administration at P1, Q/+ mice were placed under 24-hour video monitoring at P21 and P30, and seizures above Racine score 4 were counted ^18^. On P21, the control ASO group had 21 seizures (n = 8 mice), while only one seizure was observed in the Scn2a ASO ED_50_ group (n = 16 mice). The Scn2a ASO ED_80_ group was seizure free on P21 (n = 10 mice). On P30, one seizure was observed in both Scn2a ASO ED_50_ (n = 17 mice) and Scn2a ASO ED_80_ group (n = 13 mice). Because not all seizures have behavioural manifestation, 24-hour ECoG recordings were performed (Fig. 3d). No electrical seizures were detected in Scn2a ASO ED_50_ Q/+ mice during P40-45 (Fig 3e, g). Furthermore, the number of spikes were similar to age matched untreated +/+ mice (Fig. 3f). When the ECoG was repeated during P56-78, inter-ictal spikes were significantly increased, and seizures were detected (Fig. 3e-g). It was also found that Scn2a mRNA level returned to 87.6 %, 79 days after Scn2a ED_50_ administration (Fig. 3h), suggesting that seizure re-occurrence is correlated with re-appearance of Scn2a expression.

### Behavioural phenotype of Scn2a ASO treated Q/+ mice

Current anti-epileptic drugs poorly address the developmental impairment seen in SCN2A early seizure onset DEE, thus a battery of behavioural tests was performed to examine the disease modifying capability of Scn2a ASO beyond seizure control (Fig. 4a). The Q/+ mice administered with Scn2a ASO at P1, regardless of dose, had similar body weight at P30 when compared to age matched untreated +/+ mice (Fig. 4b). Because motor movement disorders were reported from patients with DEE caused by SCN2A loss of function DNMs ^14^, the motor function of Scn2a ASO treated Q/+ mice was first examined. The Scn2a ASO ED_80_ treated Q/+ mice performed similarly to +/+ mice on grip strength and grid walk test (Fig. 4c, d). However, they displayed higher ambulatory time in the locomotor assay (Fig. 4e). Increased duration in the open arm of elevated plus maze was previously reported in Scn2a haploinsufficient mouse models ^19-21^. Similarly, Scn2a ASO ED_80_ Q/+ mice spent more time in the open arm on the elevated plus maze (Fig. 4f). SCN2A null variants have been implicated in autism spectrum disorder, therefore social function was also examined ^12^. In the three-chamber social interaction test, Scn2a ASO ED_80_ treated Q/+ mice spent similar time interacting with a novel mouse as +/+ mice, indicating normal social approach (Fig. 4g). When the Scn2a ASO dose was lowered from ED_80_ to ED_50_, the altered behaviour observed in the locomotor assay and elevated plus maze was abolished, indicating that these behavioural changes could be rectified by dose titration.

**Figure 4.**
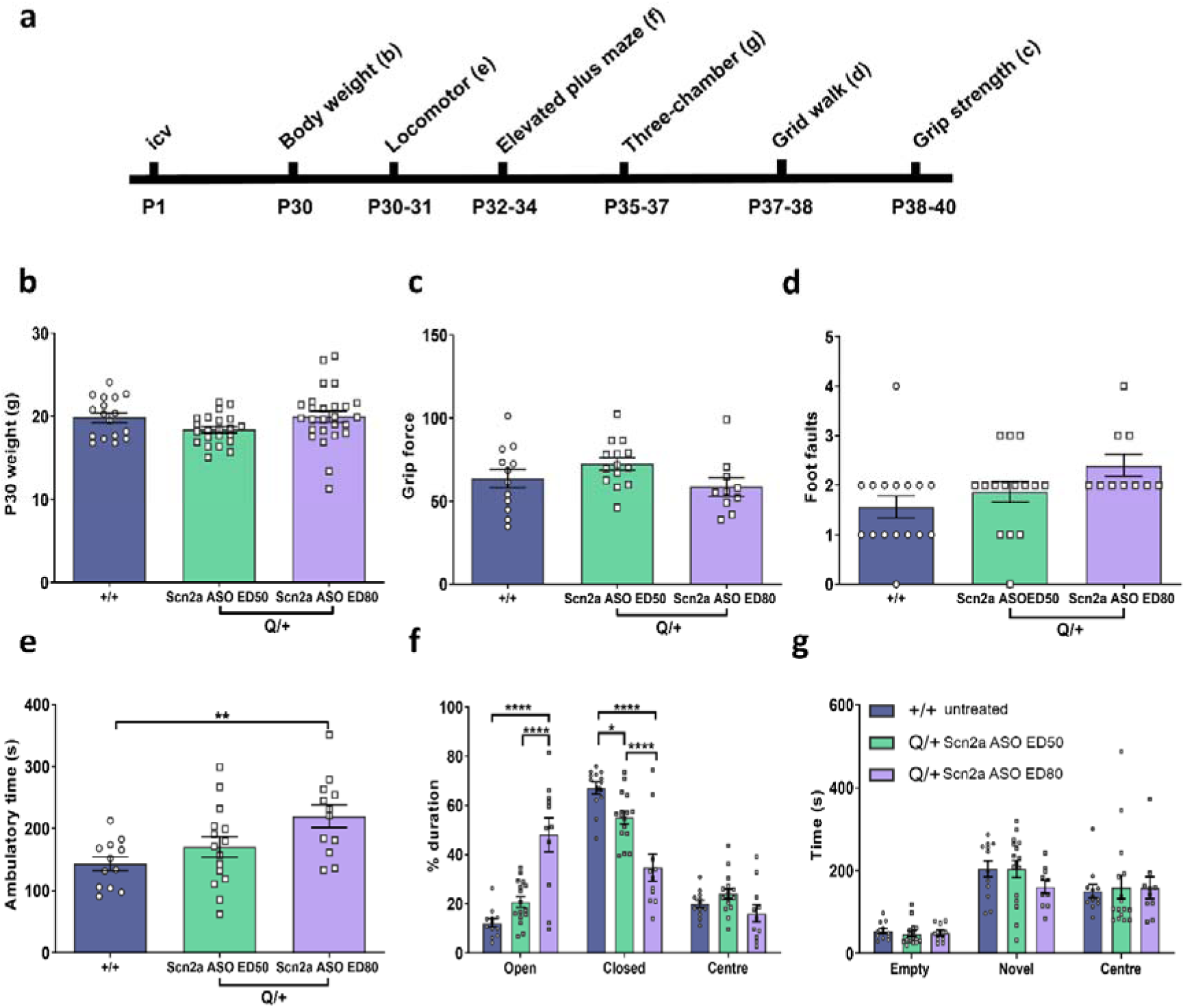
Behavioural comparison between Scn2a ASO treated Q/+ mice and untreated +/+ mice. **a.** Timeline for behavioural studies. **b.** Body weight measured on P30. **c.** Grip force. **d.** Number of foot faults in grid walk test. **e.** Ambulatory time measured in locomotor chamber. **f.** Duration spent in different arms of the elevated plus maze. **g.** Time spent in the different compartments of three chamber social interaction test. N = 12-16 per treatment group for all behavioural studies. * *P* < 0.05, ** *P* < 0.001, **** *P* < 0.0001, one-way ANOVA with Dunnett’s multiple comparison (**b-e**), two-way ANOVA with Tukey’s multiple comparison (**f-g**).

### Scn2a ASO normalises action potential firing frequency of Q/+ pyramidal neurons

To elucidate the cellular mechanism underlying Scn2a ASO efficacy, whole-cell recording on somatosensory cortical layer 2/3 pyramidal neurons was performed. In comparison to control ASO treated +/+ mice, pyramidal neurons from control ASO treated Q/+ mice showed overt gain-of-function phenotype including higher action potential (AP) frequency (AP) (Fig. 5a, b), lower rheobase (Fig. 5c), and higher input resistance (Fig. 5d). When treated with Scn2a ASO ED_80_, the AP firing frequency, rheobase and input resistance of Q/+ mice were all reverted to +/+ level (Fig. 5a-d), suggesting reduced firing of pyramidal neurons as one of the cellular mechanisms underlying Scn2a ASO efficacy.

**Figure 5.**
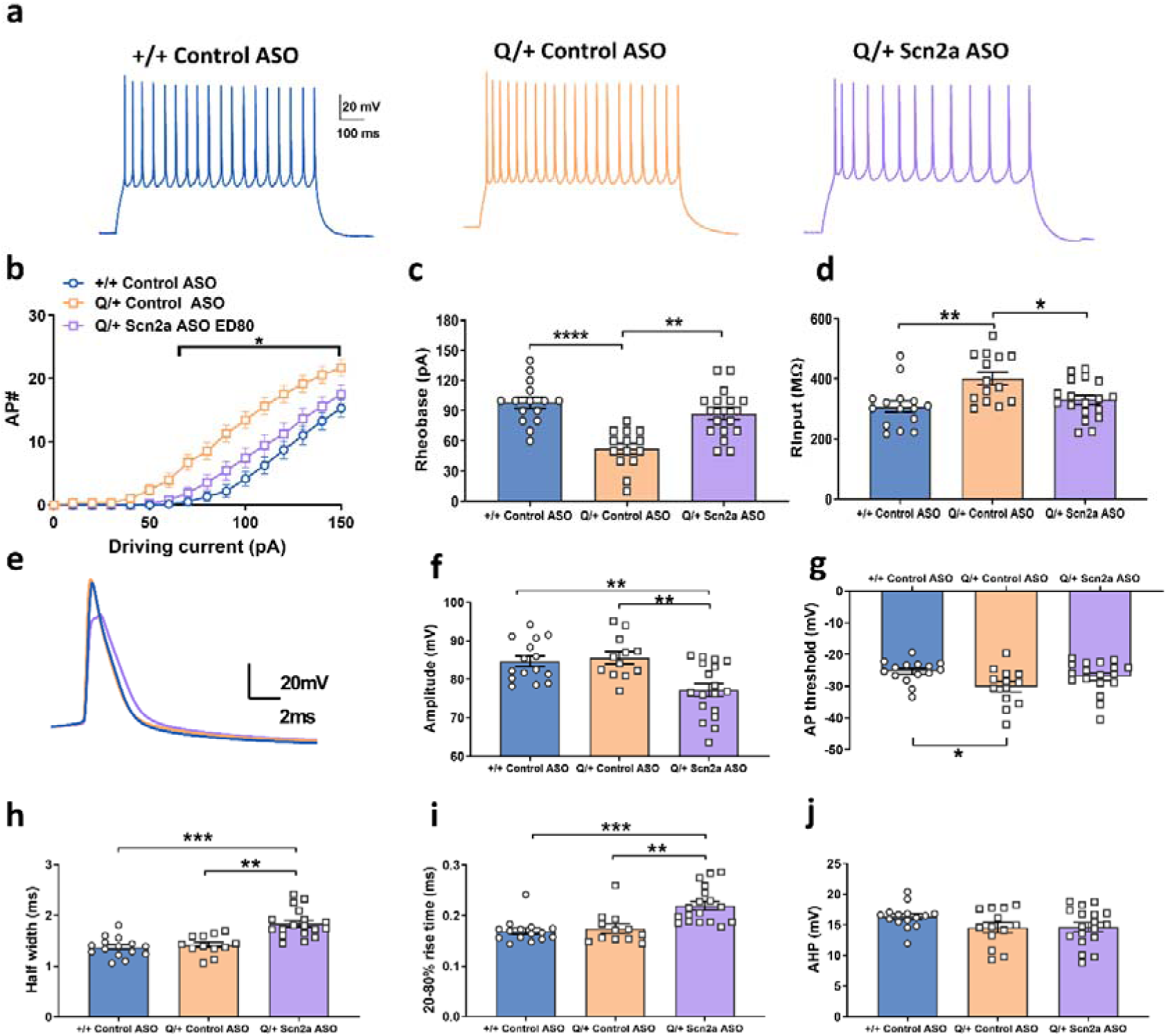
Effect of Scn2a ASO on L2/3 pyramidal neuron action potential firing. **a.** Representative voltage traces from a neuron injected with 150 pA current. Scale bar applies to all traces. **b**. Input-output relationship generated for each injected current step. **c.** Rheobase **d.** Input resistance. **e.** Averaged trace of first action potential fired for each treatment group. f. Action potential threshold **g.** Action potential amplitude **h.** Half width i. Rise time (20-80 %) j. After-hyperpolarizing potential amplitude. Control ASO +/+ (n = 15), control ASO Q/+ (n = 12), Scn2a ASO ED_80_ Q/+ (n = 18). * *P* < 0.05, ** *P* < 0.005, *** *P* < 0.001, **** *P* < 0.0001, two-way ANOVA with Tukey’s multiple comparison **(b)**, one-way ANOVA with Dunnett’s multiple comparison (**c-d, f-j**).

Examination of the first AP fired found neurons from control ASO treated Q/+ mice had significantly more hyperpolarized threshold than control ASO treated +/+ mice but the AP threshold was similar between control ASO treated +/+ and Scn2a ASO treated Q/+ mice (Fig. 5e, f). It was also found that the first AP fired from Scn2a ASO Q/+ expressing neurons had smaller amplitude, wider half width and slower rise time when compared to both control ASO +/+ and Q/+ neurons (Fig. 5g-i). No significant change was detected in the after-hyperpolarizing potential between genotype or treatment groups (Fig. 5j). However, when examining action potential fired later (tenth AP fired 50 pA after rheobase), control ASO Q/+ neurons had smaller after-hyperpolarizing potential when compared to control ASO +/+, and this was rectified by Scn2a ASO treatment (Extended data fig. 4d). The AP fired later from Scn2a ASO Q/+ neurons still had lower amplitude, slower rise time and wider half width compared to control ASO +/+ and Q/+ neurons (Extended data fig. 4a-c). This altered AP waveform was similar to that reported in an Scn2a haploinsufficient mouse model ^22^.

**Extended data figure 4.**
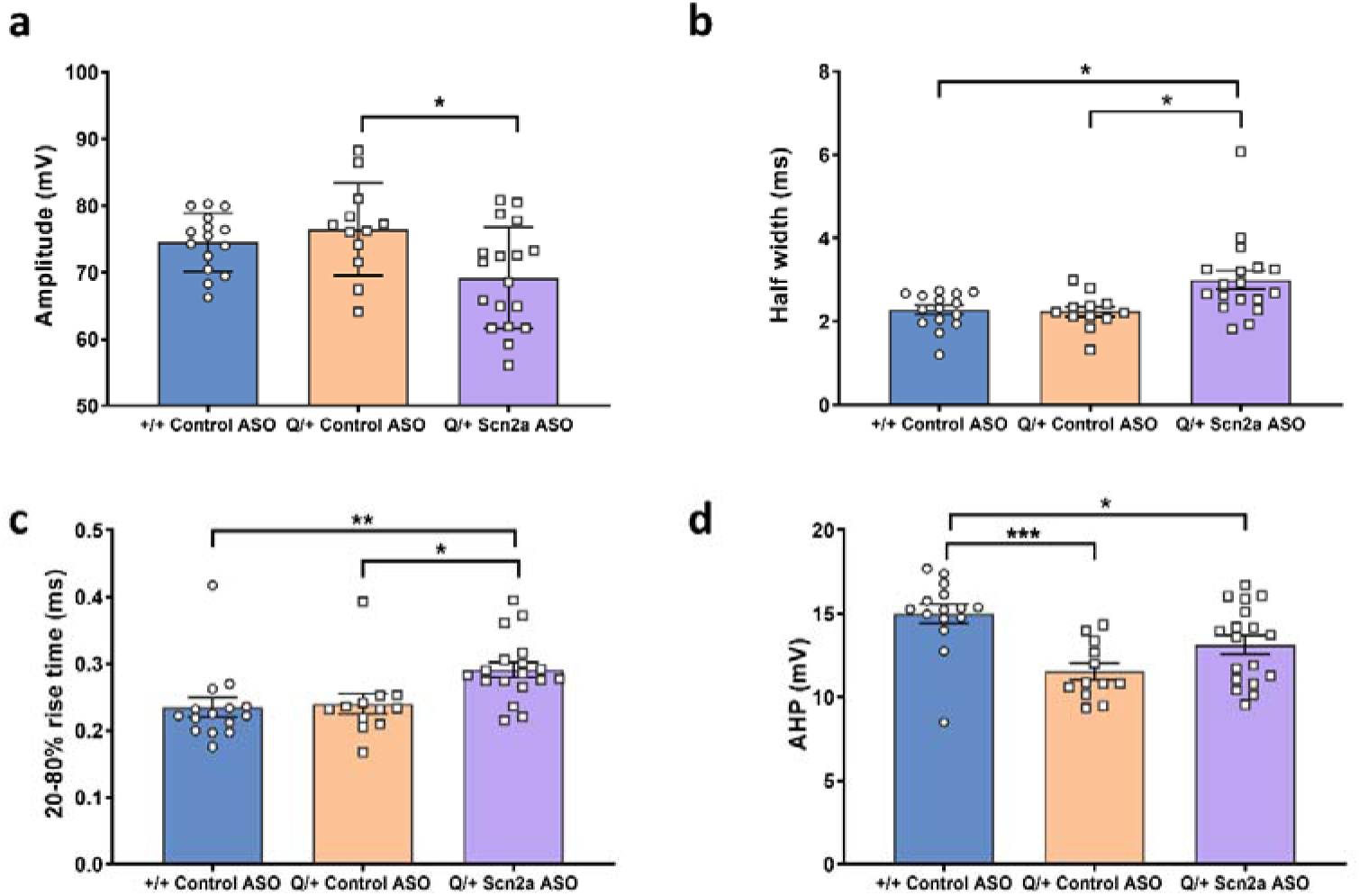
Morphology of the tenth action potential fired 50 pA after rheobase. **a.** Action potential amplitude **b.** Half width **c.** Rise time (20-80 %) **d.** After-hyperpolarizing potential amplitude. control ASO +/+ (n = 15), control ASO Q/+ (n = 12), Scn2a ASO ED_80_ Q/+ (n = 18). * *P* < 0.05, ** *P* < 0.005, *** *P* < 0.001, one-way ANOVA with Dunnett’s multiple comparison.

### Tolerability of Scn2a ASO in neonatal +/+ mice

Tolerability of Scn2a ASO was examined in +/+ mice as part of the pre-clinical safety assessment. It is known that Scn2a depletion in early rodent development is detrimental and Scn2a homozygous knockout mice die perinatally ^22,23^. Consistently, administration of Scn2a ASO at ED_80_ in P1 +/+ mice resulted 43 % lethality between P11-25 but survival then stabilised to the end of experiment at P80 (Fig. 6a). This is likely to be because Scn8a becomes the predominant voltage gated sodium channel isoform in the rodent brain after P30 ^10^. The Scn2a ASO ED_80_ +/+ mice were smaller in size, with a significantly lower body weight compared to control ASO +/+ mice (Fig. 6b, d). In contrast, Scn2a ASO ED_50_ was not lethal in neonatal +/+ mice (Fig. 6a), and the body weight was similar to control ASO treated +/+ mice (Fig 6b, d).

**Figure 6.**
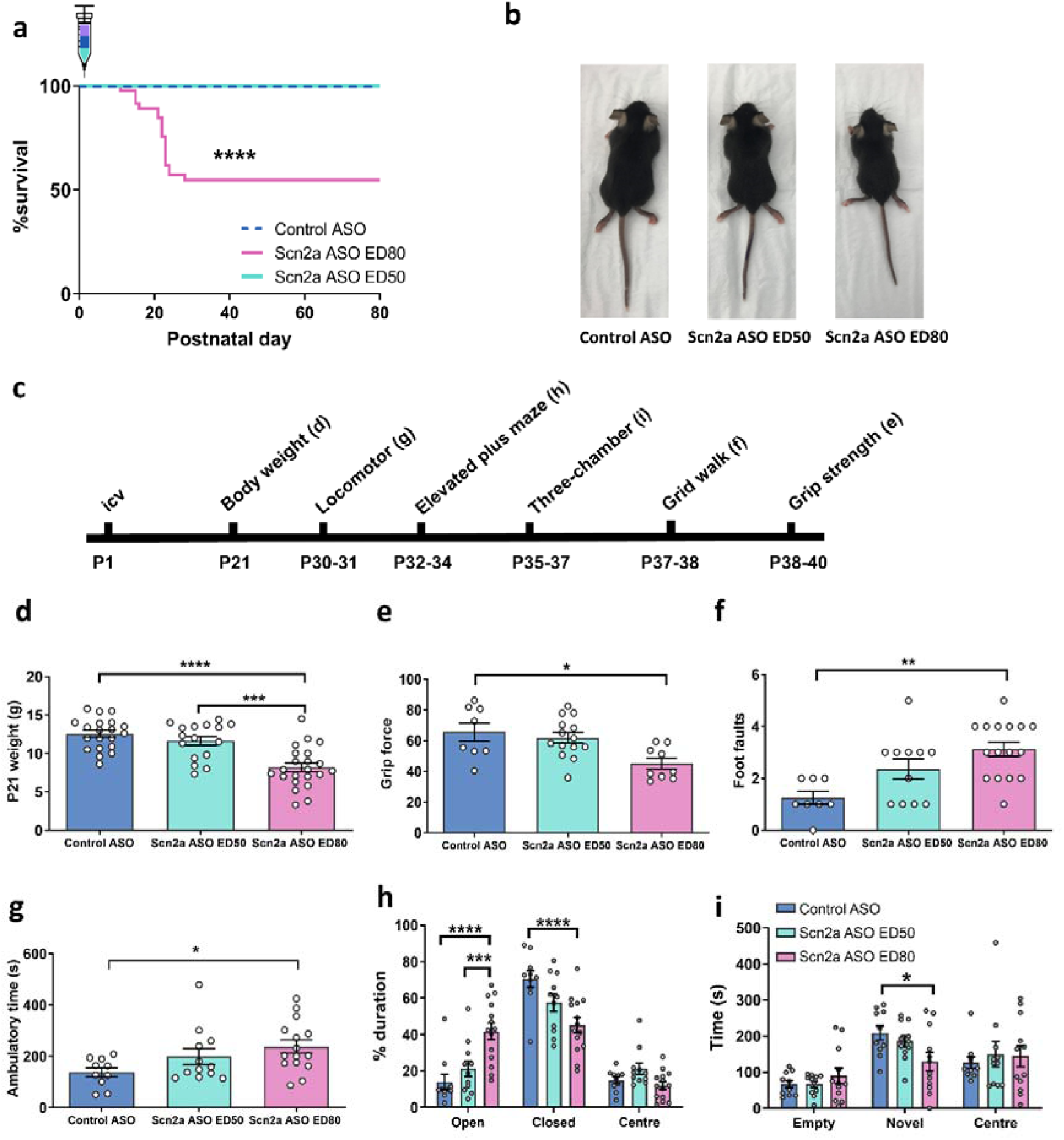
Effect of Scn2a ASO administration in P1 +/+mice. **a.** Survival curves of +/+ mice administered with control ASO or Scn2a ASO on P1. control ASO (50 µg, n = 32), Scn2a ASO ED_80_ (10 µg, n = 47), Scn2a ASO ED_50_ (2 µg, n = 15). **b.** Photo illustrating the difference in body size. **c.** Timeline for behavioural studies. **d.** Body weight measured on P21. **e.** Grip force. **f.** Number of foot faults in grid walk test. **g.** Ambulatory time measured in locomotor chamber. **h.** Percentage duration spent in different arms of the elevated plus maze. Time spent in the different compartments of three chamber social interaction test. N = 8-16 per treatment group for all behavioural studies. * *P* < 0.05, *** *P* < 0.001, **** *P* < 0.0001, Log-rank test (**b**), one-way ANOVA with Dunnett’s multiple comparison (**d-g**), two-way ANOVA with Tukey’s multiple comparison (**h**-i).

Behavioural studies were performed on +/+ mice administered with control ASO or Scn2a ASO at P1 (Fig. 6c). Scn2a ASO ED_80_ reduced grip strength (Fig. 6e), which could be linked to their smaller body size. It also caused higher number of foot faults on the grid walk test, indicating a motor learning deficit (Fig. 6f). Similar to Q/+ mice, Scn2a ASO ED_80_ +/+ mice showed higher ambulatory time in the locomotor assay and spent more time in the open arm on elevated plus maze (Fig. 6 g, h). This confirmed that the altered behaviour observed in Scn2a ASO ED_80_ treated Q/+ mice was unrelated to disease. Furthermore, ASO ED_80_ +/+ mice spent less time interacting with novel mice in the three-chamber test, indicating a social deficit (Fig 6i). When the dose of Scn2a ASO was lowered to ED_50_ +/+ mice showed no behavioural deficits were observed across all tests performed.

### Exaggerated pharmacology of Scn2a ASO in adult +/+ mice

Since the role of Scn2a is predominantly replaced by Scn8a later in development ^10^, it was hypothesised that the effect of Scn2a ASO overdose will be less pronounced when administered to adult +/+ mice (P49). Adult +/+ mice were overdosed with Scn2a ASO that caused more than 80 % Scn2a mRNA reduction (300 µg or 500 µg, Extended data fig. 5a). Despite the overdose, no lethality was observed, indicating Scn2a is not critical for survival later in development. This suggests the therapeutic window of Scn2a ASO widens with age. Furthermore, +/+ mice that received Scn2a ASO on P49 performed similarly to control +/+ mice on locomotor assay, Y-maze, acoustic startle test and paired-pulse inhibition test (Extended data fig. 5b-e). However, Scn2a ASO overdose led to decreased interaction with novel mice in three-chamber test, the inability to recognise novel object or food, elevated anxiety in dark-light test, and poorer performance on rota-rod when compared to control +/+ mice (Extended data fig. 5g-j). This indicates that while Scn2a ASO overdose in adult +/+ mice is not lethal, it may still affect motor and cognitive traits in rodents.

**Extended data figure 5.**
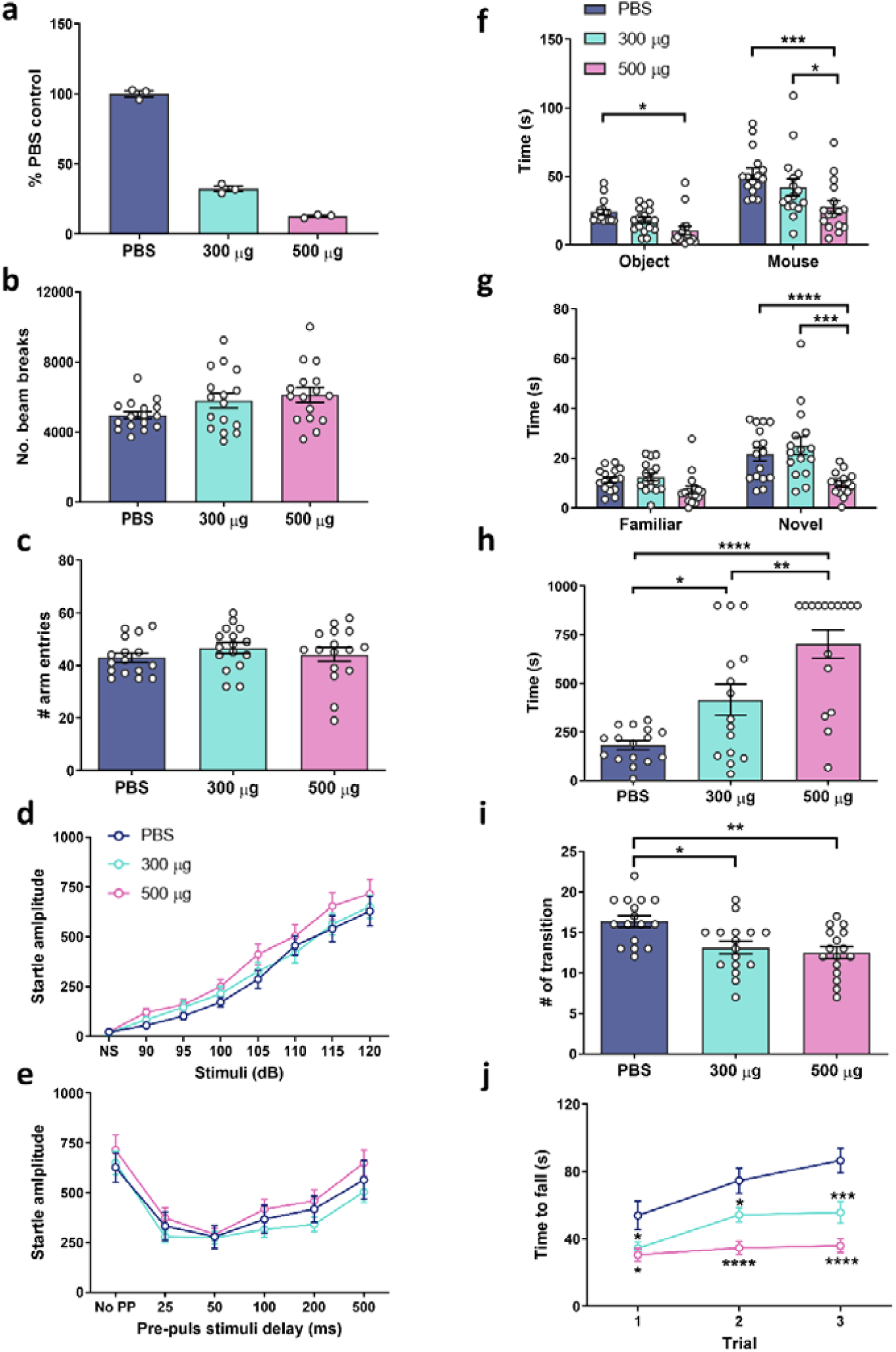
Behaviour of +/+ mice icv injected with Scn2a ASO on P49. **a.** Scn2a mRNA reduction 4 weeks post icv injection. **b.** Number of beam breaks on locomotion assay. **c.** Number of arm entries in Y-maze. **d.** Startle amplitude in response to 86 dB pre-pulse stimuli delay. **e.** Startle amplitude in response to a range of acoustic stimuli. **f.** Time spent with novel object or novel mouse in three-chamber test. **g.** Time spent with familiar or novel object. **h.** Duration to locate unfamiliar food. **i.** Number of zone transitions in light dark test. **j.** Time to fall measured in rota-rod test. N = 16 mice per treatment group for behavioural studies. * *P* < 0.05, ** *P* < 0.01, *** *P* < 0.001, **** *P* < 0.0001, two-way ANOVA with Tukey’s multiple comparison (**d-g**, j), one-way ANOVA with Dunnett’s multiple comparison (**b-c, h**-i).

## Discussion

In this study we have shown that antisense oligonucleotide targeting Scn2a successfully and specifically reduced levels of mRNA and protein expression, which provided therapeutic benefits in a mouse model carrying one of the most recurrent SCN2A gain-of-function disease variants. Scn2a ASO treatment prolonged the life span in these mice and rescued the seizure phenotype. Behavioural analysis revealed that the treated mice had a phenotype comparable to the wild type mouse. With further testing of different Scn2a ASO concentrations and injection regimens we could show that repeated application of a lower dose of ASO provided the best outcome. Moreover, injection of the ASO at a later time point was also of significant benefit. These results are pertinent as multiple dosing as well as therapy commencement after the onset of the disease are most likely scenarios expected in the clinical setting. Lastly, given the critical role of SCN2A and the fact that its loss of function has been linked to different disease phenotypes, we have importantly shown that a therapeutic dosage of ASO would not produce any adverse effects.

Following thirty years of development, ASO technology has gained a significant recognition with the recent clinical success of ASO treatments for spinal muscular atrophy, Duschenne muscular dystrophy and Huntington’s disease ^5-7^. The ASO approach has several advantages over other gene therapy approaches. Firstly, the introduced change does not affect genetic information carried in the DNA but target the intermediate RNA molecules and the resulting protein production, circumventing potential irreversible off-target changes to the patient genome. Secondly, the effect of ASO on RNA is transient and any potential adverse effects can be stopped simply by refraining from further application. Lastly, the effect is dose-dependent and can be adjusted enabling identification of a specific therapeutic window for each target and/or patient. ASOs show different modes of action which can either affect the stability of mRNA or its splicing and interactions with other molecules. Early seizure onset DEE caused by SCN2A gain-of-function variants have poor clinical outcomes and the correlation between genetic variant, functional effect and clinical phenotype is well defined^12,13,17^, making it a credible candidate to trial ASO therapy to down-regulate the expression of SCN2A.

To provide evidence that an ASO is a viable therapeutic strategy for SCN2A gain-of-function DEE, we generated a knock in mouse model (Q/+) carrying one of the most recurrent gain-of-function SCN2A variants, p.R1883Q (mouse equivalent to human p.R1882Q), linked to early seizure onset DEE. Compared to other existing Scn2a gain-of-function mouse models in the literature ^24,25^, the Q/+ mouse model has a severe disease phenotype, presenting with spontaneous seizures and premature death phenotype, with all mice dying before P30. Patch clamp recordings in slices from Q/+ mice showed left-shifted input-output curve. This increased firing rate in principal neurons is a likely cellular mechanism of disease that has also been reported in other SCN2A gain-of-function models. Altogether, these data indicate that we had generated a validated model of SCN2A gain-of-function disease.

Despite the disease severity, a single bolus of Scn2a ASO was able to dose-dependently extend survival and reduce seizure frequency of Q/+ mice. Remarkably, Scn2a ASO administration led to a seizure free period in this model. This effect was correlated with Scn2a ASO mediated Scn2a down-regulation at both mRNA and protein level. The cellular mechanism of action was also elucidated whereby Scn2a ASO reversed multiple biophysical gain-of-function features of Q/+ expressing neurons to +/+ mouse neurons’ level.

Previous studies have shown that the *in vivo* half-life of ASO is several months, which we also confirmed. The long-lasting action of ASOs indicates that only infrequent dosing may be required in patients. We have observed re-emergence of spiking, seizures and sudden death approximately two months after initial injection at P1. Importantly, re-dosing at a later time point resulted in further protection from the severe seizure phenotype. This has important implications for therapy as it suggests a similar pattern of ASO application as the one currently used for Nusinersen where upon initial “loading” subsequent injections occur every 3-4 months. Furthermore, re-emergence of pathology that can be rescued by another ASO injection indicates that the acute impact of the pathogenic variant must underlie some proportion of the DEE phenotype. Importantly, administration of Scn2a ASO two weeks after disease onset was still able to extend survival and reduce seizure frequency. In contrast, the current first line therapy, phenytoin, showed a significantly lesser disease reversing effect. It is possible that a higher phenytoin dose could lead to improved survival, however an increased dose may lead to phenytoin poisoning in neonatal mice ^26,27^.

One of the biggest challenges in DEE management is to relieve the disease burden caused by both seizure and developmental impairment ^28,29^. While existing anti-epileptic drugs can control seizures in a subset of patients with early seizure onset SCN2A DEE, developmental impairment remains unattended ^14,17^. Due to the early premature death phenotype, the behavioural profile of Q/+ mouse model could not be ascertained. Nevertheless, we could show that Q/+ mice administered with Scn2a ASO ED_50_ on P1 had a similar behavioural profile to +/+ mice on a range of motor and psychosocial tests.

Since Scn2a is critical for early postnatal development, the therapeutic window of Scn2a ASO is often limited by exaggerated pharmacology caused by action against the intended target ^30^. Scn2a is a critical gene for early rodent development, and it is known that Scn2a deficiency is lethal^23^. Consistently, Scn2a ASO ED_50_ is tolerable when administered into P1 +/+ mice, but ED_80_ resulted in 43 % lethality between P11-25. It is essential to note at this point that the lethal consequence of an exaggerated Scn2a ASO pharmacology was not found in the Q/+ mice. Furthermore, later in development when Scn8a expression increases ^10^, reducing Scn2a by more than 80% was not lethal in +/+ mice, suggesting that exaggerated pharmacology of Scn2a ASO is dependent on disease state and age of administration.

Scn2a is vital for normal development, as behavioural deficits were reported in haploinsufficient mouse models ^19,20,31,32^. Behavioural features such as hyperactivity in locomotor cell, decreased interaction with novel mice, and increased time spent on open arm on elevated plus maze were observed in both Scn2a haploinsufficient mice, and mice treated with Scn2a ASO ED_80_ on P1 ^19,20,31^. On a cellular level, the changes in action potential morphology were also similar between excitatory neurons from Scn2a ASO ED_80_ treated Q/+ mice and Scn2a haploinsufficient mice ^22^. Overall, this confirms the behavioural and cellular alterations observed with Scn2a ASO ED_80_ are related to exaggerated pharmacology, and thus can be mitigated by dose adjustment. The starting dose, age of administration and dosing frequency will be pivotal in the evaluation of a human analogue of Scn2a ASO in Good Laboratory Practice Pharmacology (GLP) toxicology studies. Results from our study suggest ED_50_ is well tolerated in mice. However, SCN2A gain-of-function diagnosis becomes critical when considering administering an ASO dose higher than ED_50_, especially during early development.

The clinical spectrum of SCN2A disorder continues to expand, and many variants implicated in late seizure onset DEE and autism spectrum disorder are loss-of-function, which would unlikely benefit from the SCN2A down-regulatory ASO ^12,33^. However, ASO technology have been shown to upregulate gene and protein levels ^34,35^, therefore presenting an alternative therapeutic strategy for disorders caused by SCN2A loss-of-function variants.

In summary, this pre-clinical study demonstrated the therapeutic viability of ASO-mediated gene silencing for *SCN2A* gain-of-function disorders. The clinical development pathway for ASOs is well established and this study provides crucial efficacy and toxicology preclinical data for the SCN2A gain-of-function epilepsies that can be readily translated into the clinical practice in the near future.

## Methods

### ASO and drug preparation

We screened a collection of ASOs designed to target various regions of the mouse Scn2a mRNA. After screening Scn2a ASOs for their ability to reduce Scn2a levels in cultured mouse cells, and for toxicity in WT mice, we tested several ASOs by delivering them through a single intracerebroventricular (ICV) injection into the brain of P1 mice and used the most effective and well-tolerated ASO for further studies. An ASO targeting mouse Scn2a (GCTCATGTTACTCCTACCCT) and a non-targeting control ASO (CCTATAGGACTATCCAGGAA) were used for the studies. Both ASOs were developed and synthesized by Ionis Pharmaceuticals. ASOs were synthesized as described ^36^ and were 20 bp in length, with five 2′-O-methoxyethyl (MOE) modified nucleotides at each end of the oligonucleotide, and ten DNA nucleotides in the centre. The backbone of the ASOs consists of a mixture of phosphorothioate (PS) and phosphodiester (PO) linkages: 1-PS, 4-PO, 10-PS, 2-PO and 2-PS (5′ to 3′). ASOs were re-constituted with sterile Ca^2+^ and Mg^2+^ free PBS (Gibco). Phenytoin (Sigma-Aldrich) was prepared to desired concentration in sterile saline and kept at 4 °C

### Quantitative gene expression analysis (RT-qPCR)

To determine the effect of Scn2a ASO on the mRNA expression of other sodium channel isoforms (Fig. 2a), brains were homogenized in 600 µl RLT using the Qiagen RNasy Kit (Qiagen, Valencia, CA) containing 1% 2-mercaptoethanol. Total RNA was purified further using a mini-RNA purification kit (Qiagen, Valencia, CA). After quantitation, the tissues were subjected to real time RT-PCR analysis. The Life Technologies ABI StepOne Plus Sequence Detection System (Applied Biosystems Inc, Carlsbad CA) was employed. Briefly, 30 µl RT-PCR reactions containing 10 µl of RNA were run with the RNeasy 96 kit reagents and the primer probe sets listed in the materials section. All real time RT-PCR reactions were run in triplicate. The expression quantities of Scn2a and Gapdh mRNA were calculated based on the arbitrary numbers assigned to standard curve made by serial dilution of the RNA samples from control animals. The expression level of Scn2a mRNA was normalized to that of Gapdh mRNA, and this was further normalized to the level measured in controls.

To determine the dose-response curves (Extended figure 3), total RNA was isolated from mouse right cortex using the Trizol reagent according to manufacturer’s protocols (Thermofisher). Contaminating genomic DNA was removed with DNAse treatment (DNA-free Reagents, Ambion/Life Technologies). Targeting primers for Scn2a and other sodium channel isoforms were listed in Supplementary Table 1. For RT-qPCR, oligo-dT primed cDNA was synthesized from 500 ng of total RNA using Murine Moloney Leukaemia Virus Reverse Transcriptase (Promega). RT-qPCR was performed on the ViiA 7 Real-Time PCR System using GoTaq qPCR master mix (Promega) according to the manufacturer’s protocols. Relative gene expression values were obtained by normalization to the reference gene RPL32 using the 2DDCt method.

**Supplementary table 1.**
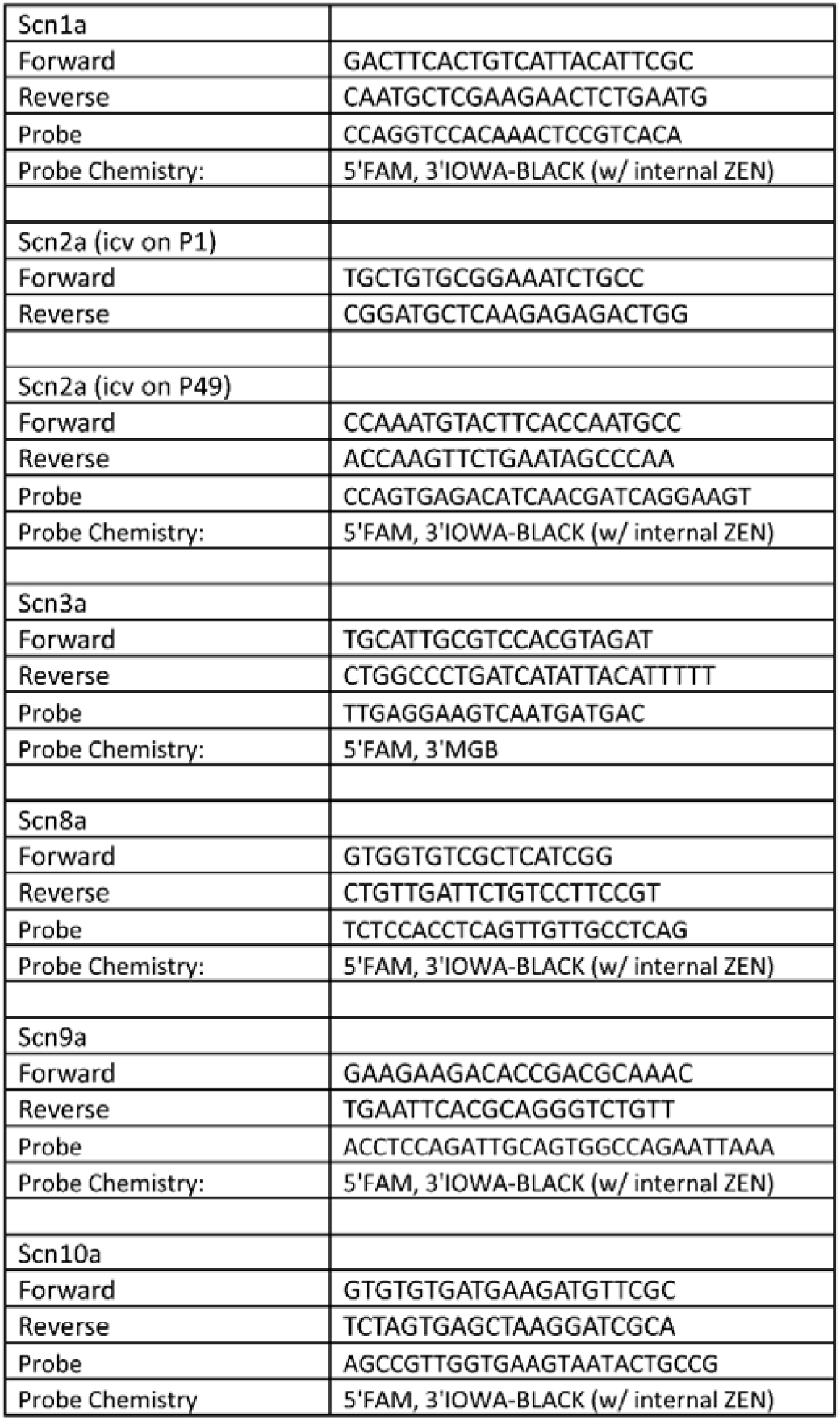
| Primers used in RT-qPCR.

### Animal model

All studies involving animals were carried out in accordance with the Guide for the Care and Use of Laboratory Animals and were approved by the Florey Institute Animal Ethics Committee. Wildtype (+/+) mice or mice heterozygous for Scn2a p.1883Q, equivalent to human mutation p.R1882Q (Q/+), were commercially generated on the C57/Bl6N background (Cyagen). Briefly, targeted embryonic stem cell clones were selected for blastocyst microinjection and implanted into surrogate dams. The F1 litters were used for experiments. All animals were maintained in a temperature-controlled room, with a 12-hour light on/off cycle and free access to food and liquid.

### Intracerebroventricular (ICV) injection

P1 mice were cryo-anaesthetised, then a Hamilton syringe (32 G Huber point) was inserted midway between lamda and right eye. The injection depth was 2 mm below skin surface and a total of 2 µl was delivered into the right ventricle. Pups were gently warmed until skin colour returned to pink before being returned to home cage.

For mice over P14, ICV injection was performed using stereotaxic apparatus (Kopf). Mice were anaesthetised with 2-4 % isoflurane and placed on frame with skull surface horizontal between lambda and bregma. A small piece of skull was removed by drilling, and an injection needle (30 G, PlasticsOne) was inserted at 0.25 mm posterior, 0.8 mm lateral to the bregma. The needle was lowered to a depth of 2.5-3 mm from brain surface. A total of 5 µl was delivered into the right ventricle using the syringe pump (KD Scientific) at a rate of 0.5 µl/s. After 2-3 minutes, the needle was slowly withdrawn, and incision was sutured.

### Immunohistochemistry

To visualise Scn2a protein, mice were ICV injected with Scn2a ASO on P1 and brain tissue was isolated between P12-15. The brains were immediately frozen with liquid nitrogen vapour and stored at -80 °C until use. Coronal sections (10 µm) were cut on a cryostat (CM1850, Leica) and mounted on Superfrost plus microscope slides (VWR). After air-drying at room temperature for 30 minutes, sections were fixed in pre-cooled acetone for 8 minutes. Sections were blocked for 1 hour in room temperature with Tris buffered saline supplemented with 3 % goat serum and 0.3 % Triton. Sections were then incubated with the following antibodies: polyclonal rabbit anti-AnkyrinG antibody (Santa Cruz) at 1:1000, and monoclonal mouse anti-Scn2a protein (Neuromab) at 1:500 overnight at 4°C. After 5 washes, sections were incubated with fluorescently labelled secondary antibodies: donkey anti-rabbit-Alexa647 at 1:800 (Invitrogen) and goat anti-mouse-Alexa488 at 1:500 (Invitrogen) for one hour at room temperature. To identify the nuclei, sections were then stained with DAPI (Sigma-Aldrich). Slides were covered with Prolong Gold Antifade (Invitrogen) and stored at -30 °C. The L2/3 layer of somatosensory cortex was examined, and confocal images captured using a Zeiss Axio 780 upright microscope, using a PL-APO 40x/NA 1.4 oil objective and according to Nyquist criterion.

For ASO visualisation, mice (P60-80) were injected with PBS or 500 µg Scn2a ASO and were perfused with 4 % paraformaldehyde 12 days post administration. The brains were isolated and sequentially equilibrated in 10% to 30% sucrose (in 0.1 M PB) at 4 °C. The brains were then embedded in OCT, frozen in chilled isopentane and stored at -80 °C until use. Sagittal sections (30 µm) were cut using a cryostat (CM1850, Leica) and mounted on positively charged slides. Sections were processed for immunohistochemistry to label neurons using (NeuN; 1:500; Millpore) and ASO backbone (1:7500; Ionis) with a guineapig and rabbit polyclonal primary antibodies respectively. Tissue was incubated with primaries diluted in 5 % normal goat serum and 0.15 % Triton-X in 0.1M PB overnight and washed several times before incubating with secondary antibodies, donkey anti-guineapig Alexa594 (Biotum) and donkey anti-rabbit Alexa 647 (Invitrogen) for 2 hours at R.T. Sections were then washed in 0.1 M PB; DAPI (1:10,000; Sigma-Aldrich) stained and mounted using ProLong Diamond antifade reagent (ThermoFisher) and cover slipped. Confocal images were acquired using a Zeiss Axio 780 upright microscope. Mosaic and z-step images were taken with a PL-APO 20x/NA0.8 air objective and high-resolution images using the PL-APO 63x/NA 1.4 oil objective and according to Nyquist criterion. All images were then deconvolved using Huygens Essential (V15.10; Scientific Volume Imaging).

### Mass spectrometry

Mouse brains were stored at -80°C after removed from animals. Approximately 100 mg of cerebellum tissue was removed and added in a 1:5 ratio to extraction buffer, which consisted of 1x Protease inhibitor, 1% sodium deoxycholate, and 100mM TEAB (Triethylamonium bicarbonate). Samples were vortexed and sonicated using the Bioruptor® Plus (Diagenode) until completely homogenized. Protein concentration was determined using the bicinchonic acid assay (BCA assay) (Thermofisher). Protein reduction was performed with DL-Dithiothreitol (DTT) to a final concentration of 5mM, for 30 minutes at 60°C. Samples were then alkylated in the dark with iodacetamide (IAA) to a final concentration of 10 mM for 30 minutes at 37°C. Samples were quenched with additional DTT to a final concentration of 10mM and incubated at 37°C for 30 minutes. The pH was checked for optimal trypsin digestion. 100µg of protein were added to Axygen Maxymum Recovery® low-bind microcentrifuge tubes, followed by the addition of trypsin in a 1:40 ratio and incubated at 37°C for 16 hours (overnight).

Deoxycholate was precipitated using 10% Formic Acid (FA). Samples were spiked with heavy peptide standard mix (JPT Peptide Technologies) and centrifuged at 12,000 x g for 10 minutes to pellet the sodium deoxycholate. The supernatant was transferred to new low-bind tubes. SPE clean up occurred using the AssayMAP Bravo Platform (Agilent) on C18 plates using the standard protocol from the manufacturer. Samples were dried down and stored in -80°C before being resuspended in 2% acetonitrile, 0.05% trifluoroacetic acid to prepare for mass spectrometry (MS).

Agilent Technologies 6495 triple quadrupole LC-MS/MS was used for the relative quantification and detection of Scn2a protein and other target peptides. The method was optimized using Skyline software (MacCoss Lab, University of Washington) and Agilent Mass Hunter software. The transitions and method settings can be seen in the supplementary material.

### Brain slice recording

Mice (P12-14) were deeply anesthetized with 4 % isoflurane, followed by brain tissue isolation. The brain was immediately transferred into ice-cold cutting solution consisting of (mM): 125 Choline-Cl, 2.5 KCl, 0.4 CaCl_2_, 6 MgCl_2_, 1.25 NaH_2_PO_4_, 26 NaHCO_3_, 20 D-glucose, saturated with carbogen (95 % oxygen and 5 % carbon dioxide). Coronal sections (300 µm) were sliced on a vibratome (VT1200, Leica). The brain slices were incubated in artificial cerebral spinal fluid consisting of (mM): 125 NaCl, 2.5 KCl, 2 CaCl_2_, 2 MgCl_2_, 1.25 NaH_2_PO_4_, 26 NaHCO_3_, 10 D-glucose saturated with carbogen for at least 1 hour at room temperature before recording.

Individual slices were placed in a recording chamber on an upright microscope (Slicescope Pro 1000, Scientifica) and perfused with aCSF at a rate of 2 ml/minute at 32 °C. Layer 2/3 excitatory neurons in the somatosensory cortex (S1) were identified with infrared-oblique illumination microscopy with a 40x objective lens (Olympus). Patch pipettes of 3-5 MΩ (Harvard Apparatus) were made using a puller (P-1000, Sutter Instruments) and were filled with internal solution consisting of (mM): 125 K-gluconate, 5 KCl, 2 MgCl_2_, 10 HEPES, 4 ATP-Mg, 0.3 GTP-Na, 10 phosphocretine, 10 EGTA and pH to 7.3 with an osmolarity of 280 mOsm. Whole-cell recording was made in current clamp mode using Axon Multiclamp 700B amplifiers (Molecular Devices). Data was acquired using pClamp v.10 software (Molecular Devices). Sampling frequency was 100 kHz and low pass Bessel filtered at 10 kHz (Digidata 1550, Molecular Devices). A holding current was injected to maintain membrane potential at approximately -70 mV. Neuronal excitability was determined by measuring voltage during a series of 800 ms steps from -60 to +300 pA in 10 pA increments every 2 s. Data was analysed using the Axograph X software (Axograph). Action potentials were determined using a 50 mV/ms threshold. Morphology was analysed on the first AP fired.

### Spontaneous seizure monitoring

Q/+ mice were placed under 24-hour video monitoring on P21 and P30 in their home cage using a digital camera (EVO2, Pacific Communications). Videos were reviewed by two experimenters and seizures over Racine score 4 were noted 18. The averaged group seizure was calculated by dividing the total number of seizures in a treatment group over the number of mice.

### Electrocorticography (ECoG)

Mice (P30-35) were anaesthetised with 2-4 % isoflurane and placed in a stereotaxic apparatus (Kopf) with the skull surface horizontal between lambda and bregma. Epidural electrodes, 1 reference and 2 recording (Pinnacle Technology), were placed on the brain surface. A ball of silver wire was used as a ground electrode. All electrodes were connected to a headmount (Pinnacle Technology) and secured to the skull with dental cement. Mice were allowed to recover for 5-7 days before ECoG recording. Brain cortical activity was sampled at 250 Hz for 24 hours (Pinnacle Technology), with a low pass and high pass filter of 40 Hz and 0.5 Hz respectively. During recording, mice could move freely in the cage with food and water ad libitum. ECoG data was analysed using ClampFit 10.7 (Molecular Devices), and a spike is identified when the amplitude was 2.5 times of baseline and duration was shorter than 80 ms.

### Behavioural Tests

All behavioural experiments were performed during the light phase between 10:00 and 17:00 hour, with a minimum of overnight recovery between tests. Mice were acclimatized to the testing rooms for at least 30 minutes before experimentation. All equipment was thoroughly cleaned and disinfected with 70-80 % ethanol between trials or testing of each mouse.

### Locomotor test

Spontaneous locomotor activity of mice ICV on P1 was measured automatically with a laser-beam system monitoring distance and movements according to prescribed parameters (MedAssociates). Each mouse was placed in the centre of the activity cage (27 × 27 cm) for 30 minutes under dimmed lighting. The ambulatory time or laser-beam breaks were determined using MedAssociates software. For mice ICV on P49, locomotor activity was measured in polycarbonate cages (22 x 20 cm) placed into frames (25.5 x 47 cm) mounted with two levels of photocell beams at 2 and 7 cm above the bottom of the cage (San Diego Instruments). Mice were tested for 120 minutes and Noldus Ethovision XT software was used to determine the number of photocell beam breaks.

### Grid walk test

Mouse was placed on an elevated grid apparatus (30 cm) and allowed to freely walk along a 1 m grid of stainless steel bars (3 mm diameter) normally spaced 1 cm apart. The test was performed after three successful training runs. For the test, the grid bars in the last 50 cm were replaced with one of several set patterns that contain missing bars at a gap on 1 or 2 bars. Behaviours on the last 50 cm grid were recorded with a digital camera (EVO2, Pacific Communications). Videos were reviewed by experimenter, and the number of foot faults (total miss or deep slip) were noted 37.

### Grip strength

A grip strength meter (Bioseb) was used to assess strength and control of the fore and hind paws. Mice were rested on the angled stainless-steel mesh assembly, grasping the mesh with all four limbs. Mice were gently pulled by the tail away from the mesh, and the maximum force prior to release of the paws was recorded. Four trials were performed on each mouse, performed by two experimenters.

### Elevated plus maze

The base of the maze was custom made of Perspex containing two open arms (7 x 31 cm) and two enclosed arms (7 x 31 x 15 cm) extending from a central platform (5 x 5cm) and was elevated 40 cm above the floor. At the beginning of the experiment, each mouse was placed in the centre of the maze facing the enclosed arm. Trial duration was 10 minutes. The duration on the open and closed arms of elevated plus maze was tracked by the TopScan software (Clever Sys Inc).

### Three chamber social interaction test

Mice were subjected to three chamber social test to assess social interaction behaviours. For mice ICV on P1, the three-chamber apparatus consisted of a Plexiglas rectangular box (40 x 38 cm) divided into three compartments with two transparent walls. The two openings on the walls allowed free access into each of the three chambers. A wired cage of equal size (16 x 10 cm) was placed in each of the two outside compartments (16 x 38 cm). The interaction zone was defined as 2.5 cm in front of the wired cage. Movement and duration were measured by the TopScan software (Clever Sys Inc.). During habituation (5 -10 minutes), the mouse was placed in the middle chamber and could freely explore all three chambers. An unfamiliar, age, strain and gender matched novel mouse was placed in the wired cage on the non-preferred side, as determined from habituation. Activity of the test subject was then monitored for a further 10 minutes. For mice ICV on P49, the three-chambered Plexiglas box (20 cm x 40.5 cm) have clear dividing walls with small semicircular openings (3.5 cm radius) allowing access into each chamber. The middle chamber is empty, and the two outer chambers contain small, round wire cages during testing. The mice are habituated to the entire apparatus for 5 min minutes. To assess sociability, mice will be returned to the middle chamber with a novel mouse (unfamiliar, strain, age and gender matched) in one of the wire cages in an outer compartment and another identical wire cage in the opposite compartment. Time spent in the chamber with the novel mouse and time spent in the chamber with the novel object was recorded for 5 minutes.

### Y-maze

The Y-maze apparatus consists of three 34 × 8 × 14 cm arms. Each mouse was tested in a single 5-minute trial and spontaneous alternations, sets of three unique arm choices, were recorded.

### Acoustic startle and pre-pulse inhibition (PPI) of the acoustic startle response

Startle and PPI testing were performed using San Diego Instruments startle chambers (SR-Lab). These consist of non-restrictive Plexiglas cylinders 5 cm in diameter resting on a Plexiglas platform in a ventilated chamber. High frequency speakers mounted 33 cm above the cylinders produce all acoustic stimuli, which are controlled by SR-LAB software. Piezoelectric accelerometers mounted under the cylinders transduce movements of the animal, which are digitized and stored by an interface and computer assembly. Beginning at startling stimulus onset, 65 consecutive 1 ms readings are recorded to obtain the peak amplitude of the animal’s startle response. A 40 min test session starting with a 10 min stimulus-free period will be used, in which pulse values are 90, 95, 100, 105, 110, 115 and 120 dB; and pre-pulse intensities are 78 dB, 82 dB and 86 dB, on a 70-dB background level with a 100 ms interstimulus interval; and 4 120 dB pulses starting the session and ending the session in order to examine habituation of the startle response. Startle pulses will be 40 ms in duration and pre-pulses will be 20 ms in duration. All trial types (pulse alone, no-stimulus trials (background only), and prepulse + pulse trials) will be presented several times in a pseudorandom order (Barros et al., 2009).

### Novel objection recognition

Mice were individually habituated to a 50 x 50 x 39 cm open field for 5 minutes. Mice were tested with two identical objects placed in the field (either two 250 ml amber bottles or two clear plastic cylinders 6 x 6 x 16cm half filled with glass marbles). An individual animal was allowed to explore for 5 minutes now with the objects present. After three trials (each separated by 1 minute in a holding cage), the mouse was tested in the object novelty recognition test in which a novel object replaced one of the familiar objects (for example, an amber bottle if the cylinders were initially used). Behavior was video recorded and then scored for contacts (touching with nose or nose pointing at object and within 0.5 cm of object).

### Olfaction test

One day before test day, an unfamiliar food high in carbohydrates (Froot Loops, Kellogg Co.) was placed overnight in the home cages of the subject mice. On the next day, each mouse was placed in cage containing 3 cm deep cedar chip bedding and allowed to explore for five minutes. The mouse was then removed from the cage, and one Froot Loop was buried in the cage bedding. The mouse was then returned to the cage and given 15 minutes to locate the buried food. Latency to find the Froot Loop and latency to initiate eating were recorded.

### Light dark test

The apparatus consisted of a rectangular box made of Plexiglas divided by a partition into two environments. One compartment (14.5 x 27 cm) is dark (8-16 lux) and the other compartment (28.5 x 27 cm) is highly illuminated (400-600 lux) by a 60 W light source located above it. The compartments are connected by an opening (7.5 x 7.5 cm) located at floor level in the centre of the partition. A mouse was placed in the dark compartment to start the 5-minute test and the time spent in each compartment was measured.

### Rota-rod

A Rota-rod Series 8 apparatus (IITC Life Sciences) was used which would record test results when the animal drops onto the individual sensing platforms below the rotating rod. An accelerating test strategy was used whereby the rod starts at 0 rotations per minute (rpm) and then accelerated at 10 rpm. Each mouse was subjected to the test 3 times per day for 3 consecutive days.

### Data analysis

All data are presented as mean ± s.e.m. Dose-response curves were fitted with data points were fitted with Motulsky regression. All statistical analyses were performed by GraphPad Prism 7, and significance determined when P < 0.05.

## References

1. Scheffer, I.E., et al. ILAE classification of the epilepsies: Position paper of the ILAE Commission for Classification and Terminology. Epilepsia 58, 512–521 (2017).

2. Maljevic, S., Reid, C.A. & Petrou, S. Models for discovery of targeted therapy in genetic epileptic encephalopathies. J Neurochem 143, 30–48 (2017).

3. Mulley, J.C., Scheffer, I.E., Petrou, S. & Berkovic, S.F. Channelopathies as a genetic cause of epilepsy. Curr Opin Neurol 16, 171–176 (2003).

4. Oyrer, J., et al. Ion Channels in Genetic Epilepsy: From Genes and Mechanisms to Disease-Targeted Therapies. Pharmacol Rev 70, 142–173 (2018).

5. Aartsma-Rus, A. & Krieg, A.M. FDA Approves Eteplirsen for Duchenne Muscular Dystrophy: The Next Chapter in the Eteplirsen Saga. Nucleic Acid Ther 27, 1–3 (2017).

6. Corey, D.R. Nusinersen, an antisense oligonucleotide drug for spinal muscular atrophy. Nat Neurosci 20, 497–499 (2017).

7. Tabrizi, S.J., et al. Targeting Huntingtin Expression in Patients with Huntington’s Disease. N Engl J Med (2019).

8. Bennett, C.F. Therapeutic Antisense Oligonucleotides Are Coming of Age. Annu Rev Med 70, 307–321 (2019).

9. Bennett, C.F. & Swayze, E.E. RNA targeting therapeutics: molecular mechanisms of antisense oligonucleotides as a therapeutic platform. Annu Rev Pharmacol Toxicol 50, 259–293 (2010).

10. Liao, Y., et al. Molecular correlates of age-dependent seizures in an inherited neonatal-infantile epilepsy. Brain 133, 1403–1414 (2010).

11. Gazina, E.V., et al. ‘Neonatal’ Nav1.2 reduces neuronal excitability and affects seizure susceptibility and behaviour. Hum Mol Genet 24, 1457–1468 (2015).

12. Wolff, M., et al. Genetic and phenotypic heterogeneity suggest therapeutic implications in SCN2A-related disorders. Brain 140, 1316–1336 (2017).

13. Sanders, S.J., et al. Progress in Understanding and Treating SCN2A-Mediated Disorders. Trends Neurosci 41, 442–456 (2018).

14. Howell, K.B., et al. SCN2A encephalopathy: A major cause of epilepsy of infancy with migrating focal seizures. Neurology 85, 958–966 (2015).

15. Carvill, G.L., et al. Targeted resequencing in epileptic encephalopathies identifies de novo mutations in CHD2 and SYNGAP1. Nat Genet 45, 825–830 (2013).

16. Allen, A.S., et al. De novo mutations in epileptic encephalopathies. Nature 501, 217–221 (2013).

17. Berecki, G., et al. Dynamic action potential clamp predicts functional separation in mild familial and severe de novo forms of SCN2A epilepsy. Proc Natl Acad Sci U S A 115, E5516–E5525 (2018).

18. Ihara, Y., et al. Retigabine, a Kv7.2/Kv7.3-Channel Opener, Attenuates Drug-Induced Seizures in Knock-In Mice Harboring Kcnq2 Mutations. PLoS One 11, e0150095 (2016).

19. Spratt, P.W.E., et al. The Autism-Associated Gene Scn2a Contributes to Dendritic Excitability and Synaptic Function in the Prefrontal Cortex. Neuron (2019).

20. Tatsukawa, T., et al. Scn2a haploinsufficient mice display a spectrum of phenotypes affecting anxiety, sociability, memory flexibility and ampakine CX516 rescues their hyperactivity. Mol Autism 10, 15 (2019).

21. Lena, I. & Mantegazza, M. NaV1.2 haploinsufficiency in Scn2a knock-out mice causes an autistic-like phenotype attenuated with age. Sci Rep 9, 12886 (2019).

22. Ogiwara, I., et al. Nav1.2 haplodeficiency in excitatory neurons causes absence-like seizures in mice. Commun Biol 1(2018).

23. Planells-Cases, R., et al. Neuronal death and perinatal lethality in voltage-gated sodium channel alpha(II)-deficient mice. Biophys J 78, 2878–2891 (2000).

24. Kearney, J.A., et al. A gain-of-function mutation in the sodium channel gene Scn2a results in seizures and behavioral abnormalities. Neuroscience 102, 307–317 (2001).

25. Schattling, B., et al. Activity of NaV1.2 promotes neurodegeneration in an animal model of multiple sclerosis. JCI Insight 1, e89810 (2016).

26. Hatta, T., et al. Neurotoxic effects of phenytoin on postnatal mouse brain development following neonatal administration. Neurotoxicol Teratol 21, 21–28 (1999).

27. Ohmori, H., Kobayashi, T. & Yasuda, M. Neurotoxicity of phenytoin administered to newborn mice on developing cerebellum. Neurotoxicol Teratol 14, 159–165 (1992).

28. McTague, A. & Cross, J.H. Treatment of epileptic encephalopathies. CNS Drugs 27, 175–184 (2013).

29. Nariai, H., Duberstein, S. & Shinnar, S. Treatment of Epileptic Encephalopathies: Current State of the Art. J Child Neurol 33, 41–54 (2018).

30. Kornbrust, D., et al. Oligo safety working group exaggerated pharmacology subcommittee consensus document. Nucleic Acid Ther 23, 21–28 (2013).

31. Shin, W., et al. Scn2a Haploinsufficiency in Mice Suppresses Hippocampal Neuronal Excitability, Excitatory Synaptic Drive, and Long-Term Potentiation, and Spatial Learning and Memory. Front Mol Neurosci 12, 145 (2019).

32. Middleton, S.J., et al. Altered hippocampal replay is associated with memory impairment in mice heterozygous for the Scn2a gene. Nat Neurosci 21, 996–1003 (2018).

33. Ben-Shalom, R., et al. Opposing Effects on NaV1.2 Function Underlie Differences Between SCN2A Variants Observed in Individuals With Autism Spectrum Disorder or Infantile Seizures. Biol Psychiatry 82, 224–232 (2017).

34. Liang, X.H., et al. Translation efficiency of mRNAs is increased by antisense oligonucleotides targeting upstream open reading frames. Nat Biotechnol 34, 875–880 (2016).

35. Liang, X.H., et al. Antisense oligonucleotides targeting translation inhibitory elements in 5’ UTRs can selectively increase protein levels. Nucleic Acids Res 45, 9528–9546 (2017).

36. Swayze, E.E., et al. Antisense oligonucleotides containing locked nucleic acid improve potency but cause significant hepatotoxicity in animals. Nucleic Acids Res 35, 687–700 (2007).

37. Farr, T.D., Liu, L., Colwell, K.L., Whishaw, I.Q. & Metz, G.A. Bilateral alteration in stepping pattern after unilateral motor cortex injury: a new test strategy for analysis of skilled limb movements in neurological mouse models. J Neurosci Methods 153, 104–113 (2006).

